# Dissecting the clonality of I1 plasmids using ORF-based binarized structure network analysis of plasmids (OSNAp)

**DOI:** 10.1101/2021.05.11.443713

**Authors:** Masahiro Suzuki, Chihiro Norizuki, Jun-ichi Wachino, Kumiko Kawamura, Noriyuki Nagano, Yukiko Nagano, Wataru Hayashi, Kouji Kimura, Yohei Doi, Yoshichika Arakawa

## Abstract

Phylogenetic relationship of 97 I1 plasmids harboring *bla*_CTX-M_ genes encoding extended-spectrum beta-lactamase (ESBL) was analyzed using the ORF-based binarized structure network analysis of plasmids (OSNAp). The majority of plasmids carrying *bla*_CTX-M-1_ or *bla*_CTX-M-8_, ESBL genes primarily associated with domestic animals, were clonal. On the other hand, plasmids carrying *bla*_CTX-M-14_ or *bla*_CTX-M-15_, identified from both humans and domestic animals, were diverse in their contents. The findings suggest that circulation of I1 plasmids among humans and animals may contribute to their diversity.

## Introduction

Extended-spectrum β-lactamase (ESBL) genes, especially *bla*_CTX-M_, are widely disseminated among *Enterobacteriaceae* (1, 2). The *bla*_CTX-M_ genes are found on plasmids of various incompatibility groups, such as F, I1, and others (3–9). Among them, *bla*_CTX-M-15_ and *bla*_CTX-M-14_ are the major ESBL genes present in *Enterobacteriaceae* from various hosts including humans and domestic animals (10–12). On the other hand, *bla*_CTX-M-1_ and *bla*_CTX-M-8_ are mainly found in *Enterobacteriaceae* from domestic animals (13–16). Phylogenetics of plasmids provide useful information on the dissemination pattern of resistance genes across various hosts. However, practical methods to compare lineages of multiple plasmids are yet to be established and validated.

We recently developed the ORF-based binarized structure network analysis of plasmids (OSNAp), a novel, digitalized approach in analyzing phylogeny of plasmids (17). With OSNAp, plasmids are converted to binary sequences based on their gene contents, and the relationship among plasmids is visualized by the neighbor-net network. The network is drawn using splits that separate plasmids into two groups based on distances between the binary sequences (Fig. 1).

**Figure 1.**
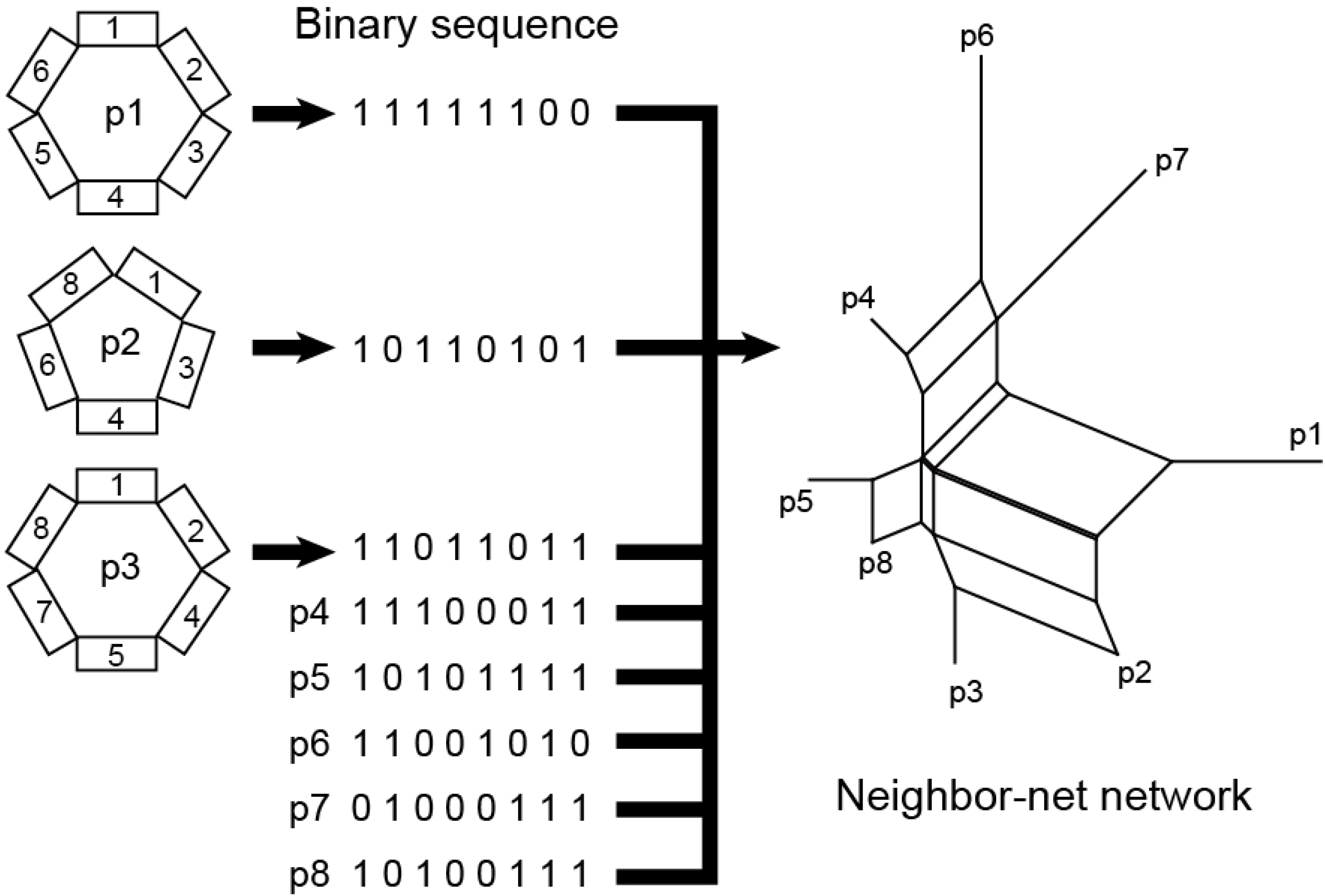
Outline of OSNAp analysis. Plasmid sequences (p1 – p3) are broken down into ORFs (1 – 8) and binarized based on the presence of ORFs. Phylogenetic relationships among plasmids are depicted as neighbor-net network based on the distance matrix of the binary sequences. ORF contents of plasmid 4 to 8 (p4 – p8) are omitted.

In this study, we analyzed *bla*_CTX-M_-carrying I1 plasmids identified in *Enterobacteriaceae* collected from multiple hosts including humans and animals. I1 plasmids were selected in this proof of concept study since their structures are relatively conserved making them suitable for OSNAp (18).

## Materials and Methods

A total of 190 I1 plasmid sequences consisting of publicly available I1 plasmid sequences (17) and I1 plasmid sequences newly obtained for this study were included. As publicly available I1 plasmid sequences, 140 I1 plasmid sequence data registered in the NCBI genome database as of 4-Aug-2018 were used (Table 1) (17). Among these publicly available plasmid sequences, 47 had *bla*_CTX-M_ genes (Table 3).

**Table 1.**
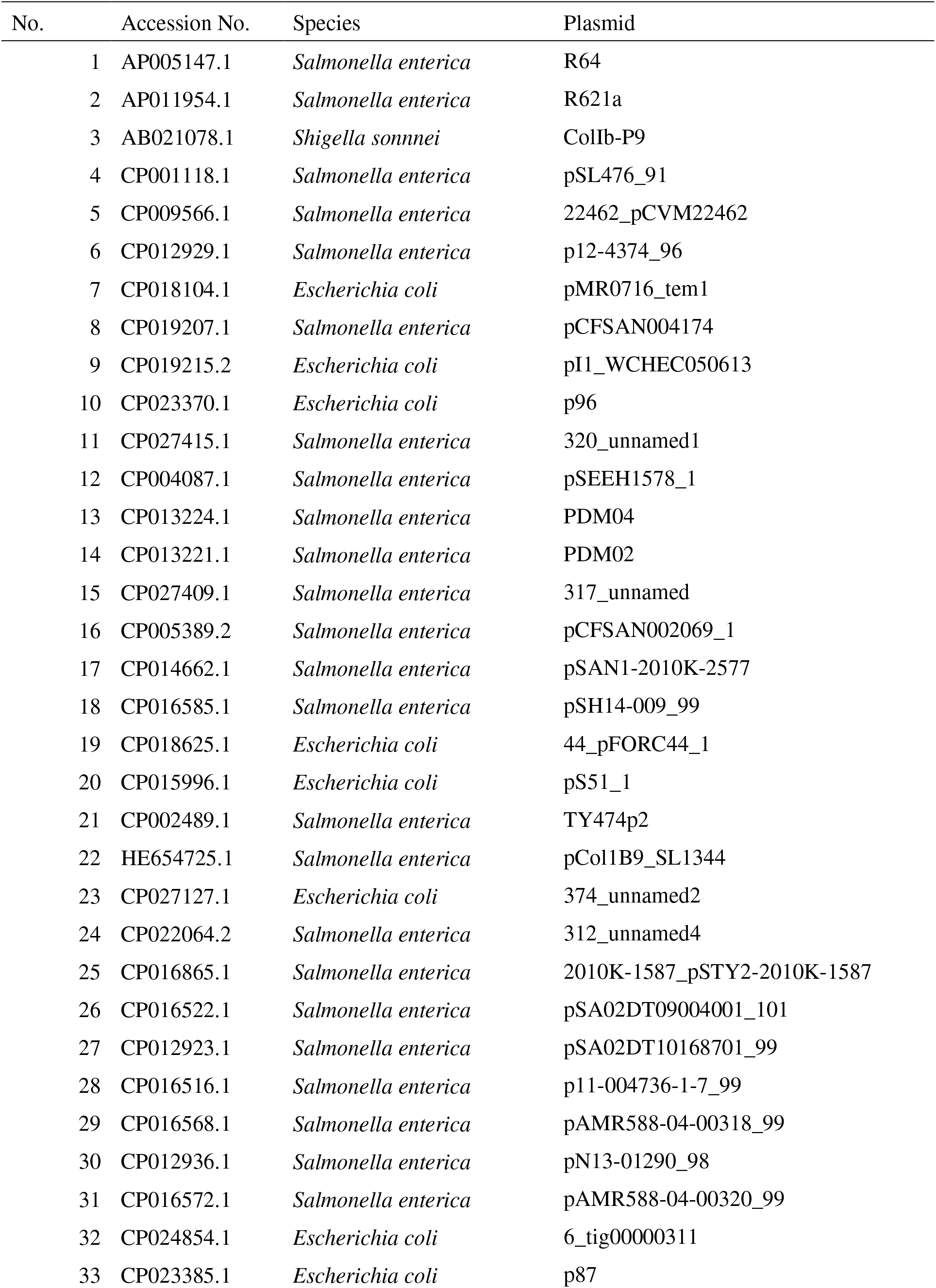
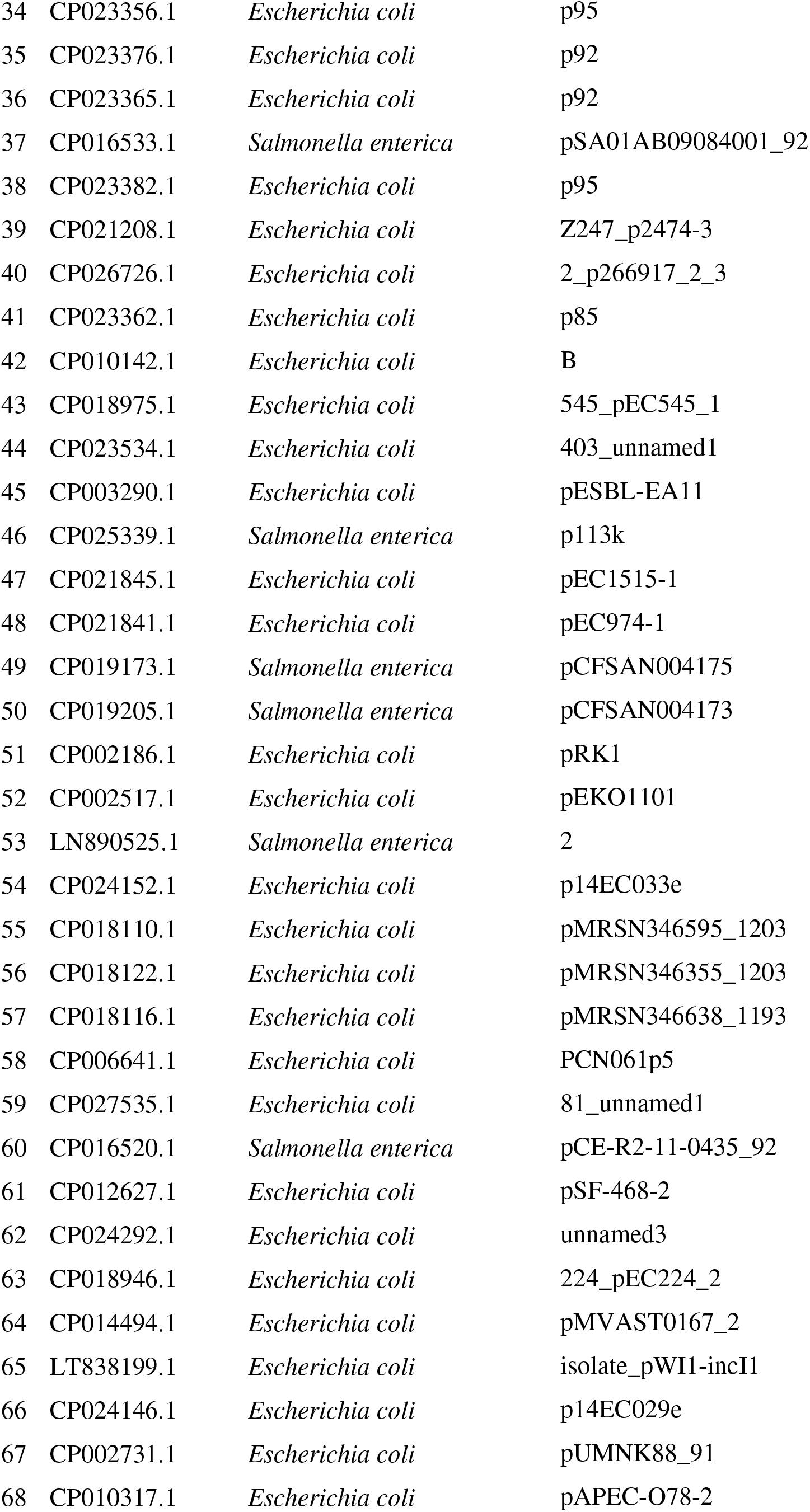
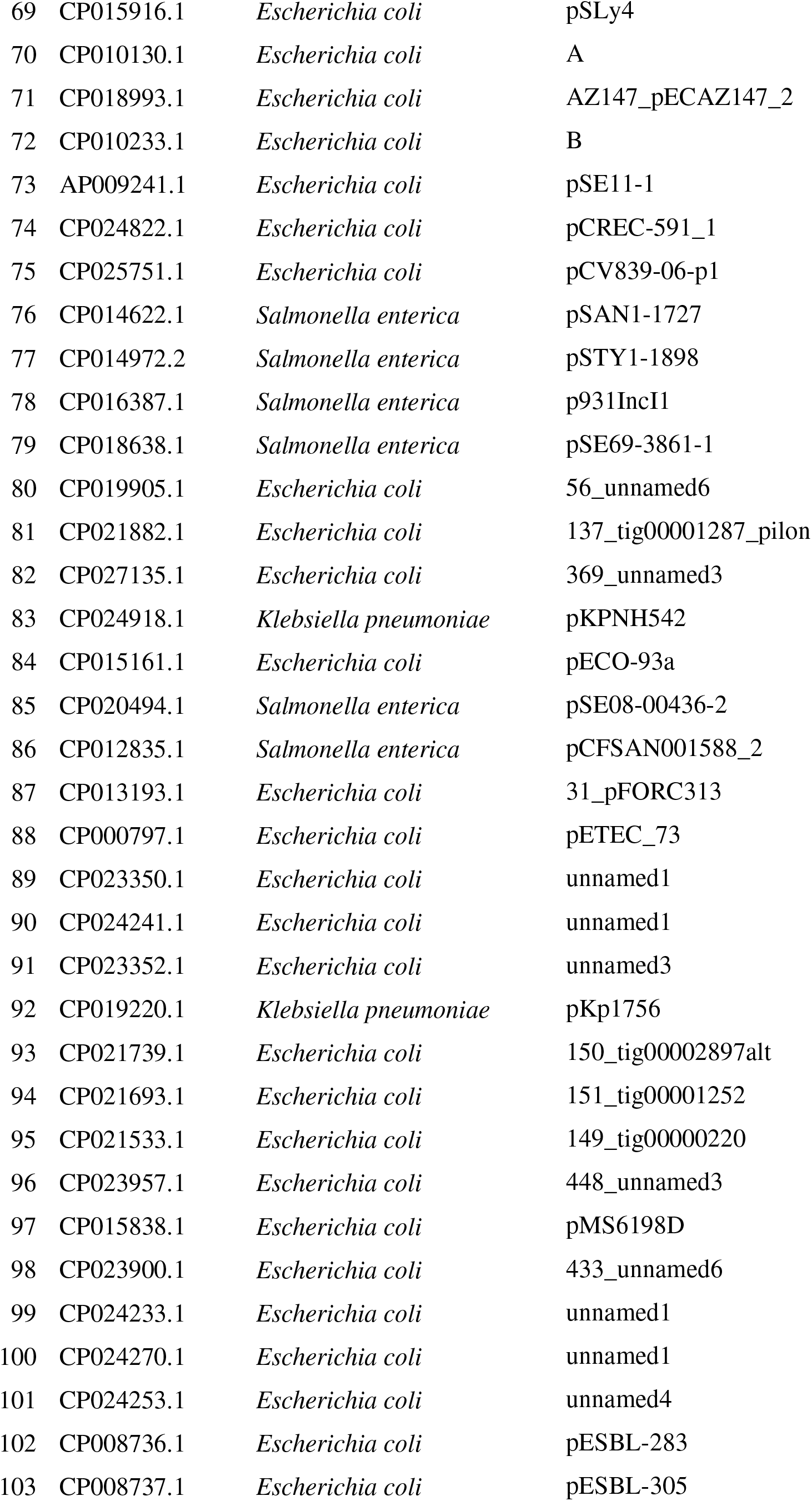
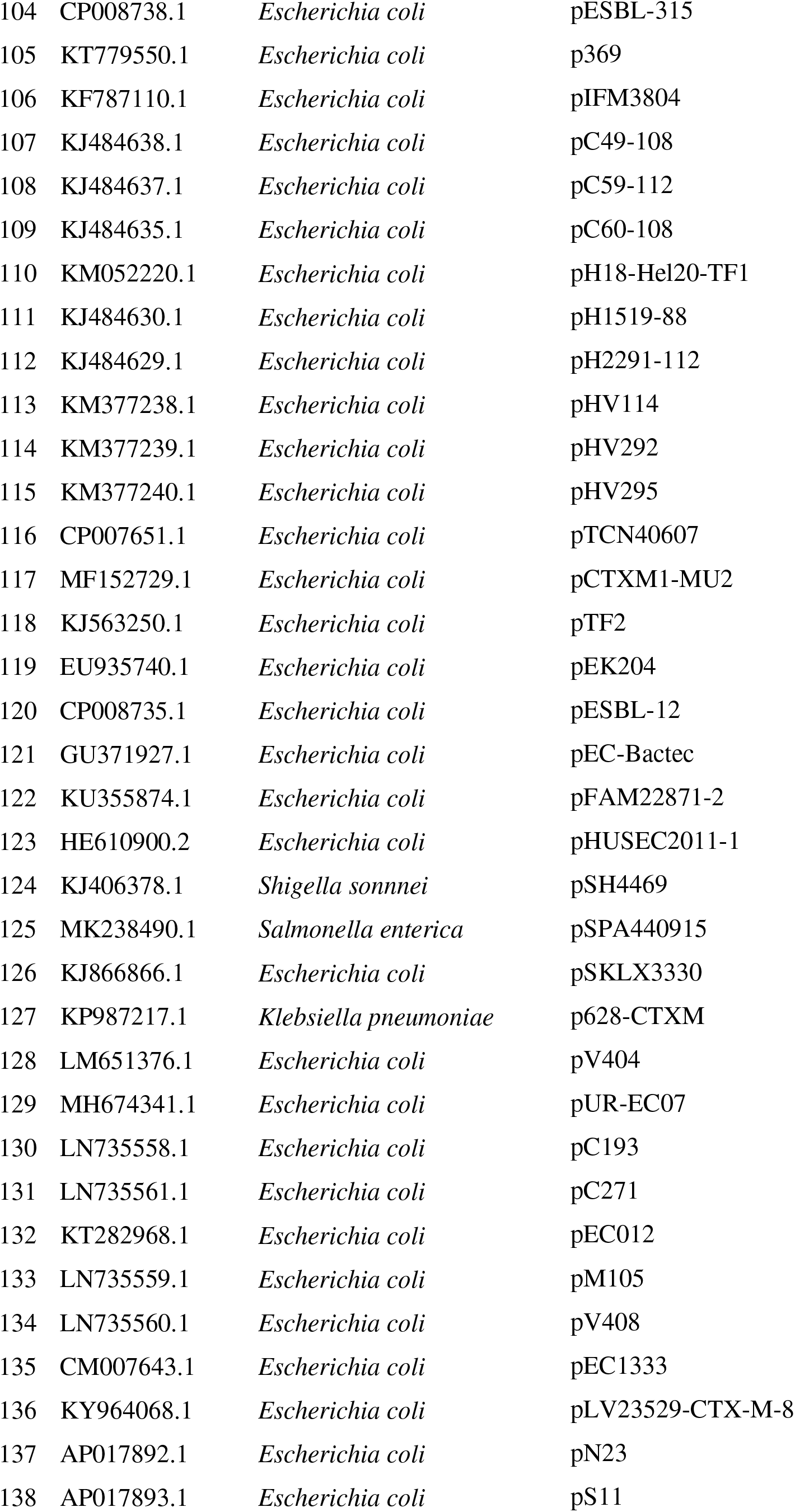
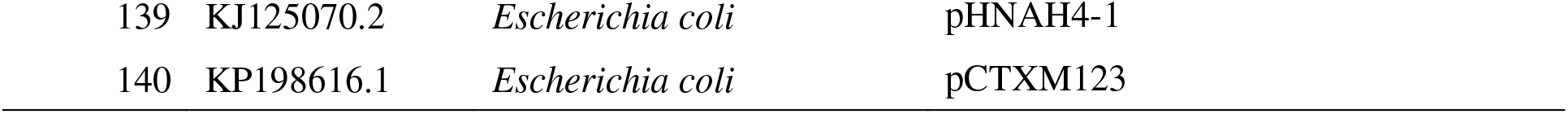
I1 plasmids obtained from NCBI database

In addition, a total of 50 I1 plasmids containing *bla*_CTX-M_ were identified in *Escherichia coli* strains isolated from humans, chicken meat, dog feces, and pig feces between 2010 to 2015. Their whole genomes were sequenced by MiSeq using QIAseq FX DNA Library Kit (Qiagen, Hulsterweg, Netherlands) and MiSeq Reagent Kit v3 600 cycles (Illumina, San Diego, CA). To enrich for sequence reads originating from plasmids, purified I1 plasmids were used as the templates for sequencing according to our previous study (8). MinION long-read sequencing was also performed according to our previous study (19). MiSeq and MinION reads were hybrid-assembled using Unicycler v. 0.4.9b, or Flye v. 2.7 and Pilon v. 1.23. Accession numbers of plasmids were listed in Table 2.

**Table 2.**
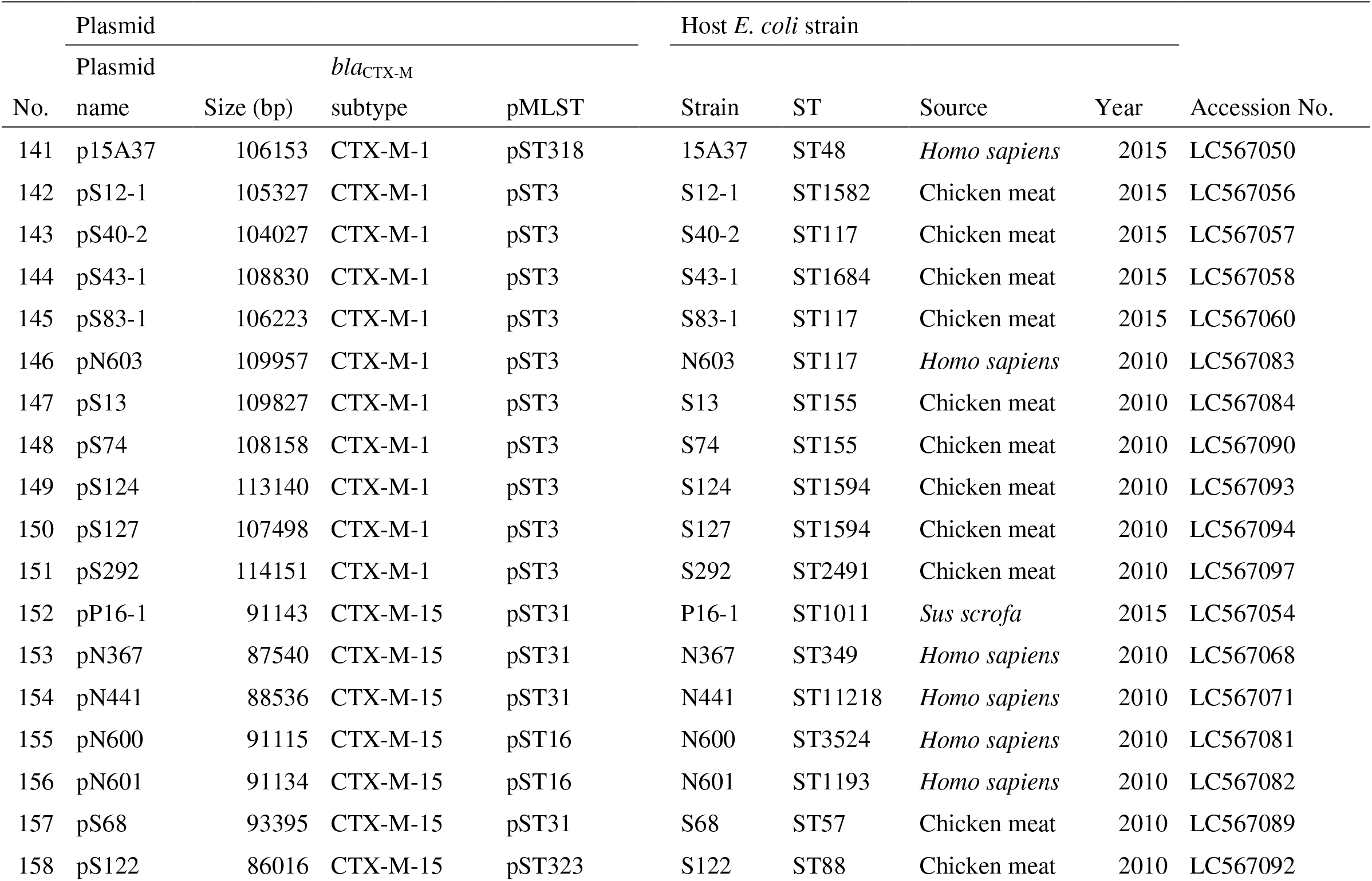
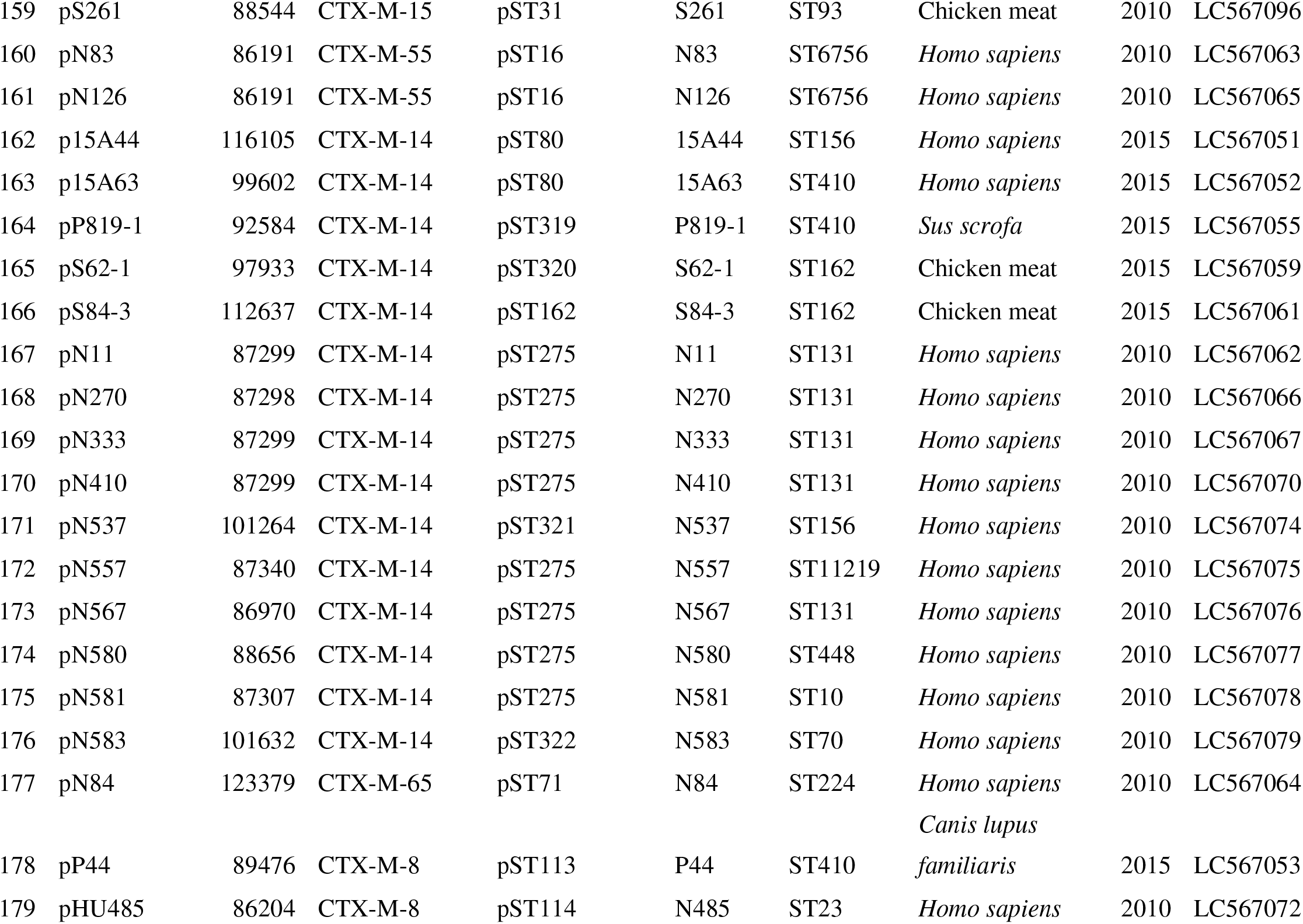
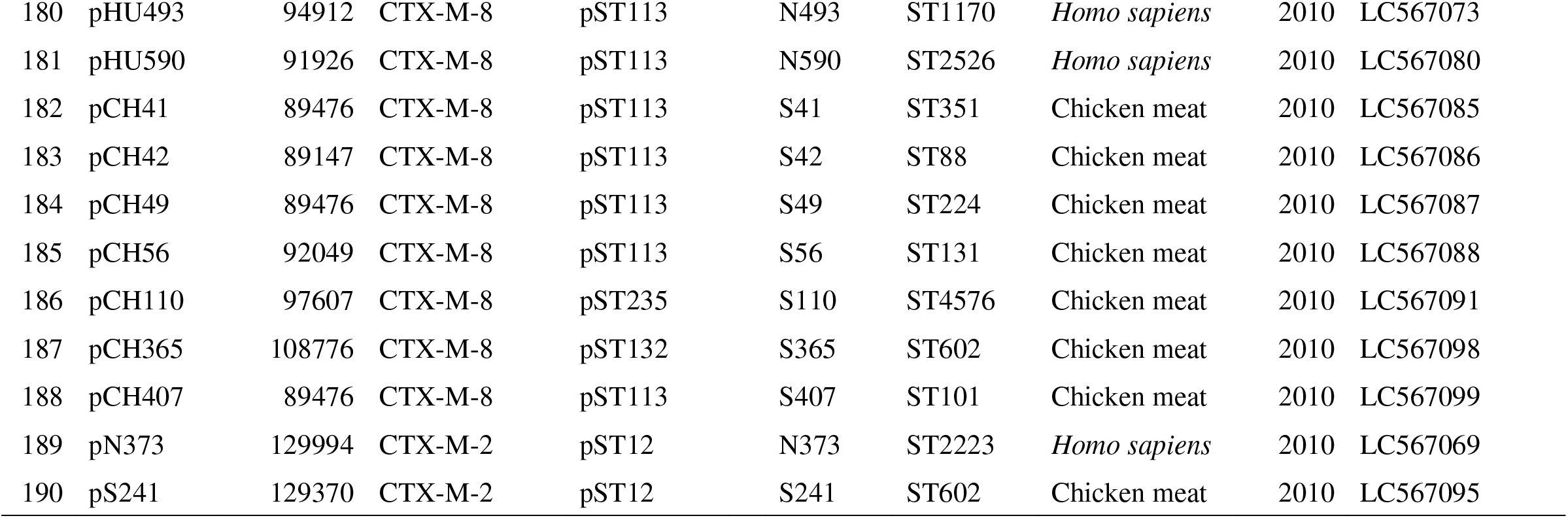
I1 plasmids analyzed in this study

**Table 3.** Meta data of I1 plasmids carrying *bla*_CTX-M_ genes

| No. | Plasmid | Source | Geographic location | Collection year | *bla*CTX-M subtype | pMLST |
| --- | --- | --- | --- | --- | --- | --- |
| 39 | Z247_p2474-3 | *Homo sapiens* | China | 2015 | CTX-M-55 | pST167 |
| 42 | B |  |  |  | CTX-M-55 | pST16 |
| 43 | 545_pEC545_1 | *Homo sapiens* | Viet Nam | 2011 | CTX-M-15 | pST16 |
| 44 | 403_unnamed1 | *Homo sapiens* |  |  | CTX-M-15 | pST31 |
| 45 | pESBL-EA11 |  |  |  | CTX-M-15 | pST31 |
| 61 | pSF-468-2 | *Homo sapiens* | USA | 2010 | CTX-M-14 | pST166 |
| 74 | pCREC-591_1 | *Homo sapiens* | South Korea | 2015 | CTX-M-123 | pST108 |
| 82 | 369_unnamed3 |  |  |  | CTX-M-15 | pSTUT01 |
| 102 | pESBL-283 | *Sus scrofa* |  |  | CTX-M-1 | pST7 |
| 103 | pESBL-305 | *Gallus gallus domesticus* |  |  | CTX-M-1 | pST3 |
| 104 | pESBL-315 | *Gallus gallus domesticus* |  |  | CTX-M-1 | pSTUnknown |
| 105 | p369 | *Gallus gallus domesticus* |  |  | CTX-M-1 | pST3 |
| 106 | pIFM3804 | *Sus scrofa* | UK | 2009 | CTX-M-1 | pST108 |
| 107 | pC49-108 | *Gallus gallus domesticus* | Switzerland | 2013 | CTX-M-1 | pST3 |
| 108 | pC59-112 | *Gallus gallus domesticus* | Switzerland | 2013 | CTX-M-1 | pST3 |
| 109 | pC60-108 | *Gallus gallus domesticus* | Switzerland | 2013 | CTX-M-1 | pST3 |
| 110 | pH18-Hel20-TF1 |  | Denmark |  | CTX-M-1 | pST49 |
| 111 | pH1519-88 | *Homo sapiens* | Switzerland | 2013 | CTX-M-1 | pST145 |
| 112 | pH2291-112 | *Homo sapiens* | Switzerland | 2013 | CTX-M-1 | pST3 |
| 113 | pHV114 | *Gallus gallus domesticus* | Switzerland | 2013 | CTX-M-1 | pST3 |
| 114 | pHV292 | *Gallus gallus domesticus* | Switzerland | 2013 | CTX-M-1 | pST3 |
| 115 | pHV295 | *Gallus gallus domesticus* | Switzerland | 2013 | CTX-M-1 | pST3 |
| 116 | pTCN40607 | *Gallus gallus domesticus* | USA | 2012 | CTX-M-1 | pST3 |
| 117 | pCTXM1-MU2 | *Sus scrofa* | Australia |  | CTX-M-1 | pST3?* |
| 118 | pTF2 |  | Denmark | 2007 | CTX-M-1 | pST49 |
| 119 | pEK204 |  |  |  | CTX-M-3 | pST16 |
| 120 | pESBL-12 | *Homo sapiens* |  |  | CTX-M-15 | pST37 |
| 121 | pEC-Bactec | *Equus caballus* | Belgium |  | CTX-M-15 | pST31 |
| 122 | pFAM22871-2 |  |  |  | CTX-M-15 | pST37 |
| 123 | pHUSEC2011-1 | *Homo sapiens* | Germany | 2011 | CTX-M-15 | pST31? |
| 124 | pSH4469 | *Homo sapiens* | Korea | 2008 | CTX-M-15 | pST16 |
| 125 | pSPA440915 | *Homo sapiens* |  |  | CTX-M-15 | pST31 |
| 126 | pSKLX3330 | *Homo sapiens* | China |  | CTX-M-55 | pST16 |
| 127 | p628-CTXM | *Homo sapiens* | China | 2010 | CTX-M-55 | pST16 |
| 128 | pV404 | *Homo sapiens* | Bolivia |  | CTX-M-14 | pST107 |
| 129 | pUR-EC07 | *Homo sapiens* | Uruguay |  | CTX-M-14 | pST80 |
| 130 | pC193 |  | Bolivia | 2011 | CTX-M-65 | pST71 |
| 131 | pC271 |  | Bolivia | 2011 | CTX-M-65 | pST71 |
| 132 | pEC012 | *Gallus gallus domesticus* | China |  | CTX-M-65 | pST71 |
| 133 | pM105 | *Homo sapiens* | Bolivia | 2011 | CTX-M-65 | pST71 |
| 134 | pV408 | *Homo sapiens* | Bolivia | 2011 | CTX-M-65 | pST71 |
| 135 | pEC1333 | sewage | Brazil | 2009 | CTX-M-8 | pST113 |
| 136 | pLV23529-CTX-M-8 | *Sus scrofa* | Portuga | 2015 | CTX-M-8 | pST113 |
| 137 | pN23 | *Homo sapiens* | Japan |  | CTX-M-8 | pST113 |
| 138 | pS11 | chicken meat | Japan |  | CTX-M-8 | pST113 |
| 139 | pHNAH4-1 | *Gallus gallus domesticus* |  |  | CTX-M-123 | pST108 |
| 140 | pCTXM123 |  | China |  | CTX-M-123 | pST108 |

Plasmid sequence types (pSTs) of the plasmids were identified using the plasmid multilocus sequence typing (pMLST) web site (https://pubmlst.org/plasmid/). Subtypes of *bla*_CTX-M_ genes were determined by ABRicate (https://github.com/tseemann/abricate) using the ResFinder database. Sequence types (STs) of the host cells were also identified using mlst command (https://github.com/tseemann/mlst). A total of 190 I1 plasmid sequences were analyzed by OSNAp under the relaxed condition (17). For the purpose of this analysis, ORFs having ≥80% sequence identity and sequence coverage were considered identical. Dice coefficient values between plasmids were also calculated. Plasmid regions (core, shell and cloud regions) were defined according to our previous study (17).

Clusters constituting the *bla*_CTX-M_ plasmids were defined by splits which maximized the number of plasmids harboring the same *bla*_CTX-M_ gene and minimized others. While clusters are identical to splits in the OSNAp analysis, the term cluster is used in this study for ease of reading. The diversity of plasmids within the same cluster was defined by the average of Dice indices.

I1 plasmid sequences were annotated using DFAST-core (https://dfast.ddbj.nig.ac.jp/dfc/distribution/) supplemented with allele-level annotation of *bla*_CTX-M_ and IS genes using ResFinder (https://bitbucket.org/genomicepidemiology/resfinder_db/src/master/) and ISfinder (https://www-is.biotoul.fr/index.php). Structures of modules carrying *bla*_CTX-M_ genes and their locations were determined from annotation data of plasmid sequences by checking the regions surrounding *bla*_CTX-M_.

## Results

### OSNAp analysis

Plasmids harboring the same *bla*_CTX-M_ gene belonged to one or two clusters, and each plasmid cluster included the same *bla*_CTX-M_, except for one cluster which contained *bla*_CTX-M-3_, *bla*_CTX-M-15_, and *bla*_CTX-M-55_-harboring plasmids (Fig. 2).

**Figure 2.**
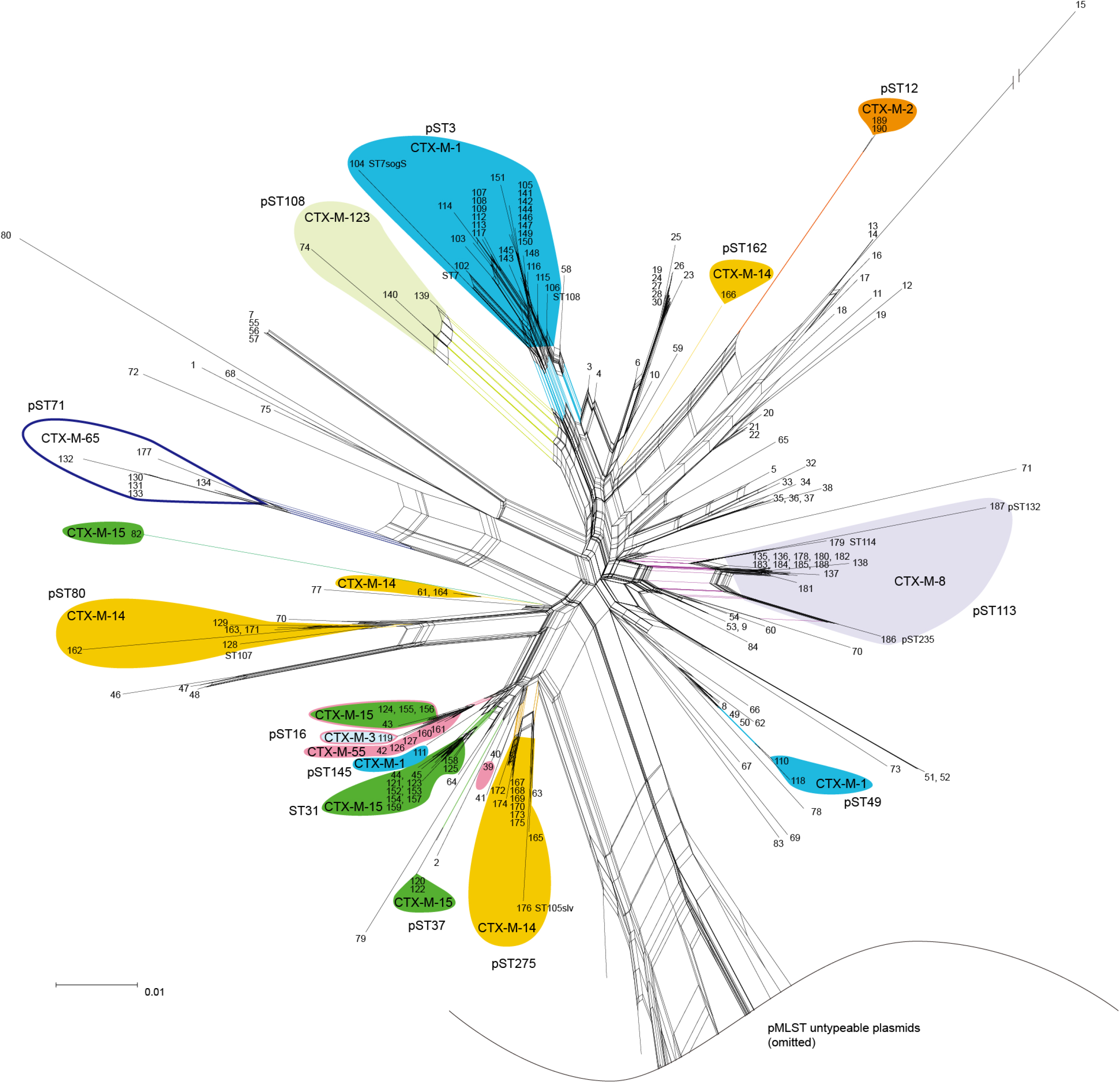
OSNAp analysis of I1 plasmids. Plasmids harboring *bla*_CTX-M-8_ were classified into single cluster and most *bla*_CTX-M-1_ harboring plasmids also belonged to a cluster. On the other hand, plasmids harboring *bla*_CTX-M-14_ and *bla*_CTX-M-15_ were classified into multiple clusters. Representative pST is shown for each cluster.

Most (27/30) plasmids harboring *bla*_CTX-M-1_ were classified into the same OSNAp cluster. The average Dice index of the *bla*_CTX-M-1_ cluster was 0.942 ± 0.036. Most plasmids in the cluster belonged to pST3 except for 3 plasmids which were of pST108, pST7, and a pST7 variant lacking *sogS*.

All 15 plasmids harboring *bla*_CTX-M-8_ were classified into the same cluster. Among them, 12 plasmids were similar with each other with an average Dice index of 0.979 ± 0.015 and belonged to pST113. However, 3 plasmids belonging to pST114, pST132 and pST235 showed lower levels of similarity compared with the other 12 plasmids. The average Dice index between the former 12 and the latter 3 plasmids was 0.868 ± 0.038.

Plasmids harboring *bla*_CTX-M-15_ were classified into 2 major and 1 minor clusters, and one singleton. The cluster including 11 *bla*_CTX-M-15_ plasmids showed an average Dice index of 0.970 ± 0.013 and represented pST31. Another major cluster also included plasmids harboring *bla*_CTX-M-3_ and *bla*_CTX-M-55_ as well as *bla*_CTX-M-15_. The *bla*_CTX-M-3_ and *bla*_CTX-M-55_ modules showed structures specific to each *bla*_CTX-M_ gene. The average Dice index of the cluster was 0.967 ± 0.016. All plasmids in this cluster belonged to pST16. The minor cluster was placed near the two major clusters in the OSNAp. Plasmids of the minor cluster belonged to pST37.

Unlike the other *bla*_CTX-M_ genes, plasmids harboring *bla*_CTX-M-14_ showed a polyclonal distribution. They were classified into 2 major and 1 minor clusters and a singleton. The largest cluster contained 10 plasmids with an average Dice index of 0.950 ± 0.049. Most plasmids in the largest cluster belonged to pST275.

### Structure of modules carrying *bla*_CTX-M_

*bla*_CTX-M-1_ was always carried on a module consisting of IS*Ecp1*, *bla*_CTX-M-1_, and an ORF for a hypothetical protein. In 25 of the 27 plasmids, the *bla*_CTX-M-1_ module was inserted between *rci* (encoding shufflon-specific DNA recombinase) and *pilVA* genes. In the remaining two plasmids, the *bla*_CTX-M-1_ module was inserted into a cloud region. The *bla*_CTX-M-1_ module was inserted into *pilJ* in the three non-pST3 plasmids.

All *bla*_CTX-M-8_ modules consisted of IS*26*, ΔIS*10*, *bla*_CTX-M-8_, and IS*26*. The modules were inserted between *impB* and *yfbB* genes.

*bla*_CTX-M-15_ plasmids of the major clusters had the same modules consisting of Tn*2*, *bla*_CTX-M-15_, IS*Ecp1* and ΔTn*2*. Plasmids of the minor cluster belonged to pST37 and had another module consisting of *bla*_CTX-M-15_ surrounded by IS*26*. The *bla*_CTX-M-15_ modules were located in a cloud region of I1 plasmids.

*bla*_CTX-M-14_ plasmids in the largest cluster had a closely related plasmid structure and their *bla*_CTX-M-14_ modules consisted of IS*Ecp1*, an ORF encoding a hypothetical protein, *bla*_CTX-M-14_ and IS*903*. In 8 of 10 plasmids, the *bla*_CTX-M-14_ module was integrated between *traH* and *traG* genes. The remaining 2 plasmids had the same *bla*_CTX-M-14_ module, which was integrated in a cloud region. However, the *bla*_CTX-M-14_ modules outside the largest cluster, including another major cluster consisting of pST80 plasmids, consisted of IS*Ecp1*, *bla*_CTX-M-14_, and IS*903*. Some of the modules were interrupted by other IS elements or partial deletion. They were integrated into the cloud region.

#### MLST of *E. coli* strains

Most of the host *E. coli* strains belonged to diverse STs even when they harbored the same *bla*_CTX-M_ genes (Table 2). Among them, 11 strains carrying *bla*_CTX-M-1_ were classified into 7 STs and no strains carrying *bla*_CTX-M-15_ or *bla*_CTX-M-8_ showed the same ST. However, strains with I1 plasmids carrying *bla*_CTX-M-14_ gene were less diverse. In particular, 5 of 6 strains carrying pST275 and *bla*_CTX-M-14_ gene belonged to ST131.

## Discussion

I1 plasmids are found in multiple host bacterial species but especially frequently in *Escherichia coli* and *Salmonella enterica*. Therefore, I1 plasmids may contribute to exchanging of antimicrobial resistant genes between these species. Moreover, I1 plasmids are capable of hosting multiple *bla*_CTX-M_ variants (20). Plasmid structures of the I1 linage are relatively well conserved, making them good targets of OSNAp, a novel approach in analyzing phylogeny of plasmids we developed recently.

Our data indicate that I1 plasmids harboring the same *bla*_CTX-M_ genes generally spread clonally among different strains since they were classified into a limited number of OSNAp clusters whereas the host *E. coli* strains belonged to various STs. Our data support previous studies suggesting that I1 plasmids harboring the same *bla*_CTX-M_ gene shared similar plasmid structures (8, 9, 21–23).

The I1 plasmids harboring *bla*_CTX-M_ could be divided into two major groups according to their similarity. One included plasmids harboring *bla*_CTX-M-1_ and *bla*_CTX-M-8_ which showed clonal dissemination overall. The other included *bla*_CTX-M-14_ and *bla*_CTX-M-15_ plasmids which showed polyclonal dissemination consisting of two or more plasmid clones.

Interestingly, clonal plasmids carrying *bla*_CTX-M_ genes are largely restricted to animals (6, 8, 22, 24). Moreover, *bla*_CTX-M_ genes in this group are mainly associated with I1 plasmids (6, 8, 22, 25, 26), and not reported from F plasmids which are the dominant plasmid type overall in *E. coli*. Therefore, *bla*_CTX-M_ genes of this group may seldom exchange *bla*_CTX-M_ genes with other plasmids. Moreover, I1 plasmids harboring them appear to spread in *E. coli* in conjunction with international trade of meat products (27).

On the other hand, *bla*_CTX-M_ genes such as *bla*_CTX-M-14_ or *bla*_CTX-M-15_ were detected from various sources including humans and domestic animals. Moreover, *bla*_CTX-M_ genes in this group were frequently found on F plasmids in addition to I1 plasmids (3, 5, 7, 28, 29). This suggests that these *bla*_CTX-M_ genes may have transmitted between I1 and F plasmids. Adaptation to a variety of host animals and plasmids may contribute to the extensive spread of this group of *bla*_CTX-M_ genes.

In this study, we analyzed I1 plasmids as proof of concept because of their relative structural simplicity. However, F plasmids are the most common plasmid type especially carrying *bla*_CTX-M-14_ and *bla*_CTX-M-15_ genes, which are the dominant ESBL genes (28, 29). The phylogeny of F plasmids is complex due to their extensive structural diversity (19, 28, 30). Phylogenetic analysis of F plasmids will therefore be challenging and may require adjustments to OSNAp such as analyzing subgroups of plasmids based on the similarity of gene contents.

In summary, I1 plasmids harboring *bla*_CTX-M_ genes have disseminated clonally, and the diversity of plasmid gene contents harboring the same *bla*_CTX-M_ subtypes may depend on their occurrence and history of their spread. I1 plasmids carrying *bla*_CTX-M-1_ or *bla*_CTX-M-8_, which are known to spread by the global meat supply chain, showed clonal phylogeny. On the other hand, *bla*_CTX-M-14_ and *bla*_CTX-M-15_ showed polyclonal phylogeny. In particular, the latter *bla*_CTX-M_ genes are frequently reported on F plasmids (3, 5, 7, 28, 29), and it can be hypothesized that their ability to integrate into various plasmids may have contributed to their success in becoming predominant *bla*_CTX-M_ genes. The ability to integrate into various plasmids is also considered to be an important factor underpinning successful spread of certain drug resistance genes. Circulation among human and animals may help diversification of plasmids carrying *bla*_CTX-M_ genes.

## Acknowledgements

This study was supported by grants from the Food Safety Commission, Cabinet Office, Government of Japan (Research Program for Risk Assessment Study on Food Safety, no. 1504), JSPS KAKENHI Grant Number JP20K10457, and AMED under Grant Number JP20fk0108061.

